# Integration of Nanomaterials and DBS to Improve Basal Ganglia Oscillations Using Delayed Van der Pol Model

**DOI:** 10.1101/2025.04.08.647898

**Authors:** M. A. Elfouly

**Affiliations:** Independent Researcher (No Institutional Affiliation)

**Keywords:** Delay Differential Equations, Mathematical Models, Nanomaterials, Deep Brain Stimulation, Neurodynamics

## Abstract

Basal ganglia play a crucial role in motor control and the challenges posed by pathological oscillations, especially in Parkinson’s disease. Although deep brain stimulation (DBS) is effective in attenuating pathological frequencies, it often leads to side effects. This study presents a comprehensive framework for understanding and improving neural oscillation dynamics by integrating nanomaterials with DBS. Nanomaterials with their controllable frequencies through length, width, and structure could stabilize oscillations and maintain normal neural frequencies. This study introduces three mathematical models: the DBS model, the nanomaterials model, and a combined model of both, which use the Van der Pol delay model to demonstrate how the periodic force and nanomaterials collaborate to stabilize brain activity. Numerical simulations indicate that DBS by itself lowers harmful frequencies but might make the motor cortex unstable because it increases the strength of signals. In contrast, the nanomaterial models reduce the amplitude and activate the motor cortex. The combination of DBS and nanomaterials significantly improves stability, reduces pathological oscillations, and decreases hyperactivity. The use of nanomaterials must be carefully monitored to ensure that they do not suppress normal neural activity, highlighting the need for experimental validation. This study provides a promising direction for the development of personalized therapeutic technologies that balance the suppression of pathological activity with the preservation of normal neural function.

## 1. Introduction

The basal ganglia are important parts of the brain that help control voluntary movement by sorting through signals from the cerebral cortex, boosting the right signals, and blocking the wrong ones. This complex neural coordination contributes to the precise and smooth execution of movements and prevents random or unintended movements [1]. The literature indicates that the basal ganglia’s function extends beyond motor tasks to include cognitive and behavioral processes, thanks to its extensive connections with the limbic system and cerebellum. Graybiel (2008) has shown that the basal ganglia are actively involved in habit formation, motivation, decision-making, and reward processing, making them an important integrating center between motor and cognitive systems [2-3].

One of the main brain disorders linked to problems in the basal ganglia is Parkinson’s disease, caused by a decrease in dopamine production due to the loss of neurons in the substantia nigra [4]. This imbalance disrupts the balance between direct and indirect neural pathways, leading to clinical symptoms such as bradykinesia, muscle stiffness, and tremors at rest [5]. Brittain and Brown (2014) have demonstrated a direct relationship between these symptoms and hyperactivity in the beta frequency range (13–30 Hz). Hypersynchronization between neurons disrupts motor signal transmission and impairs precise motor control [6].

Deep brain stimulation (DBS) is one of the most prominent modern therapeutic approaches for reducing motor symptoms associated with Parkinson’s disease [7]. This method involves placing electrodes in certain areas of the brain, like the subthalamic nucleus, to send high-frequency electrical signals that interrupt harmful brain activity and help the motor system function better [8]. Clinical results by Lozano et al. (2019) have demonstrated the effectiveness of this technique in reducing pathological oscillations and improving motor performance [9]. However, its success depends on the precision of the stimulation settings, such as frequency, intensity, and location [10]. Studies indicate that DBS may cause side effects such as fatigue and depression, especially when used to treat movement disorders. Furthermore, we still lack a complete understanding of the precise mechanisms by which DBS functions, which complicate the process of customizing optimal treatments for patients. [11-12].

Nanomaterials have emerged as promising candidates for neurotherapeutic intervention [13-14]. This field has garnered increased attention due to the unique properties of carbon nanotubes (CNTs) and graphene, including their exceptional conductivity and specific responsiveness to neural signals [15-17]. Shabani et al. (2023) demonstrated that these materials can attenuate pathological neural oscillations and enhance the brain’s response to electrical therapy [18-19]. Kulkarni et al. (2023) found that these materials can reduce oxidative stress and facilitate the transport of drugs such as dopamine across the blood-brain barrier, making them a flexible option for treating neurological disorders [20-27].

As manufacturing technologies have advanced, nanomaterials have evolved from passive drug carriers into essential components in the development of nanochips directly implanted in the brain [28]. Transparent graphene chips have demonstrated a high ability to record and stimulate neural activity without affecting surrounding tissue [29-30]. These chips offer high recording resolution and diminish immune responses, thereby augmenting their potential for long-term therapeutic applications [31-36]. These advancements signify a substantial progression toward the creation of nanoscale neural interfaces that can monitor and modulate neural activity in real time [37-41].

Mathematical models are crucial for accurately simulating the dynamic behavior of the nervous system and understanding the interaction of these technologies with it [42-43]. Nonlinear models, exemplified by the Van der Pol oscillator, have demonstrated efficacy in examining transitions between stable and chaotic states, as evidenced by the research of Rosenblum and Pikovsky (2004) [44-47]. Integrating time delays into these models is crucial due to the non-instantaneous nature of neural signal transmission [48-50]. Evidence indicates that these delays may affect the initiation or stability of pathological oscillations [51]. Elfouly and Sohaly (2022) established that incorporating delays in the Van der Pol model enhances the precision of dynamic forecasts [52].

This theoretical study proposes a novel approach to treating motor brain disorders by incorporating nanomaterials into the neural environment. These cylindrical nanostructures possess tunable resonance frequencies determined by their length and diameter. This allows for selective responsiveness to pathological oscillations, such as those in the beta frequency range linked to Parkinson’s disease. Nanotubes work like tiny shock absorbers that can reduce unusual brain activity while keeping normal signals intact because they conduct electricity well and affect specific areas.

However, their functionality may vary in the complex biological brain environment due to ionic shifts, cellular interactions, and immune responses. To solve this problem, we suggest creating smart nanotubes that turn on only when they detect harmful brain activity, similar to how advanced closed-loop DBS systems work. These smart nanotubes selectively regulate neural activity by targeting only the harmful oscillations.

Using delayed Van der Pol equations, we develop a nonlinear mathematical model to analyze this concept. This model integrates the damping effects of nanotubes and incorporates DBS as a periodic external force. The study builds upon the model by Elfouly (2024), which effectively captured the dynamic symptoms of Parkinson’s and Huntington’s diseases. Our improved model shows how nanotubes and DBS work together in both helpful and harmful ways [53].

By changing factors like the size of the nanotubes and the intensity and frequency of the DBS, the model allows us to study joint resonance, reduce oscillations, or discover new dynamic states. The study uses methods like Hopf bifurcation and harmonic balance, along with simulations, to evaluate how these changes affect amplitude, frequency, and synchronization.

This study provides a basic plan for creating smart neurotherapeutics that use nanomaterial features along with adjustable stimulation. It provides helpful information for improving treatment plans, cutting down on expensive clinical trials, and moving forward with creating personalized neural implants that fit each patient’s specific brain activity.

## 2. Mathematical Model

This section introduces a mathematical model created to examine how nanomaterials and DBS influence unusual brain wave patterns in the basal ganglia circuits. The model was built using a delayed Van der Pol oscillator to represent how the basal ganglia, thalamus, and cortex interact, while considering the natural time delays that happen in how neurons communicate.

The model reproduces three major motor states typically associated with Parkinson’s and Huntington’s diseases:

### 1. Resting Tremor

Characterized by a pre-movement readiness of the motor cortex that receives positive feedback from the indirect pathway. A long delay in this feedback causes unwanted small movements, resulting in a resting tremor because of abnormal synchronization.

### 2. Motor excitation

More brain activity strengthens the direct pathway of excitation, boosting signals from the cortex and enhancing movement abilities. This process mimics the response to therapeutic stimulation or transient dopaminergic surges.

### 3. Suppressed activity

High activity at first is reduced because of the competition between direct (excitatory) and indirect (inhibitory) pathways, leading to a lower level of cortical excitability, which is a key sign of worsening motor function.

To evaluate therapeutic modulation under these dynamic conditions, we propose three models corresponding to different intervention strategies:

#### Model 1—Nanomaterials Only

This model considers the effect of smart nanotubes embedded in neural tissue. These nanotubes, designed as smart nanosheets capable of detecting and selectively activating local pathological oscillations, replace passive agents. Their changing behavior depends on the amount of *N* (*t*), which naturally decreases when there is no stimulation and reacts to normal activation levels. The damping term − *ηN* (*t − τ*) *x* (*t − τ*) introduces a delayed, frequency-specific inhibition of pathological activity, mimicking targeted therapeutic regulation.

#### Model 2—DBS Only

Here, deep brain stimulation is modeled as a continuous, open-loop, cyclic power *ϵcos*(*Ωt*), operating independently of the internal neural state. This idealized representation mirrors conventional deep brain stimulation protocols, where high-frequency stimulation is applied to specific targets regardless of ongoing neural oscillations.

#### Model 3—Integrated Nanomaterials and DBS

This hybrid model embodies the synergistic or competitive interaction between the two therapeutic modalities. The deep brain stimulation input provides continuous stimulation, while the nanomaterials provide a state-dependent adaptive damping response. The combination of constant external stimulation and targeted adjustments based on position enables a thorough examination of the best conditions for intervention.

The variables and parameters in these models are defined as follows:

- *x*(*t*): Oscillatory activity within the basal ganglia circuit.
- *a*_1_: Strength of the delayed feedback from the direct pathway.
- *a*_2_: Strength of the delayed feedback from the indirect pathway.
- τ_1_: Time delay associated with the signal transmission through the direct pathway.
- τ_2_: Time delay associated with the signal transmission through the indirect pathway.
- α*x* (*t*) : Inhibitory effect of the hyperdirect pathway.
- *η*: Damping intensity induced by the smart chip on pathological oscillations.
- *β*: Passive decay rate of the chip in the absence of pathological input.
- *δ*: Baseline responsiveness or intrinsic activation rate of the chip.
- ϵ*Cos*(Ω*t*): External periodic input from DBS, with amplitude *ϵ* and frequency *Ω*.
- *x*(*t*≤0)=*h*: Initial condition.

#### Model 1: Nanomaterials Effects

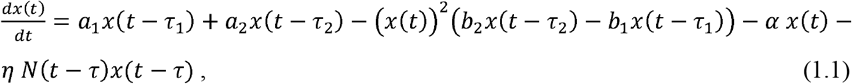

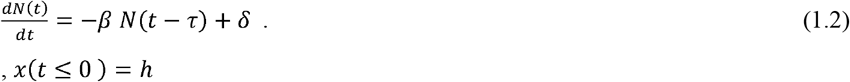

#### Model 2: DBS Effects

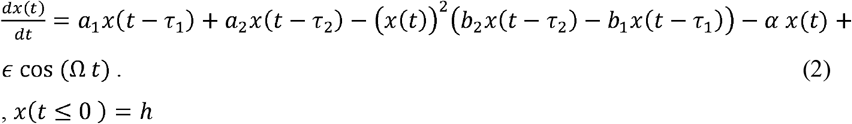

#### Model 3: Combined Effects of Nanomaterials and DBS

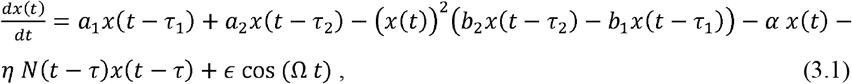

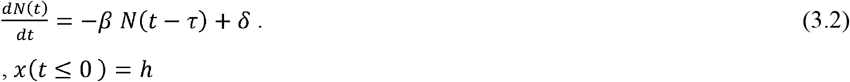

## 3. Analysis of Oscillatory Dynamics

In this section, we will derive the equilibrium points and characteristic equations from the eigenvalues of Model 1; we will use the conditions in [52] to identify the regions of stability and Hopf bifurcation between each of the two new parameters. Furthermore, we will employ the Model 2 harmonic balance method to ascertain the optimal values for the periodic solutions. We will later use these values in the numerical simulations of the three models to understand the oscillation dynamics and analyze the influence of different factors on the stability of the system.

### Model 1: Nanomaterials Effects

#### Equilibrium Points

To determine the equilibrium points, we begin by setting the derivatives in equations (1.1) and (1.2) equal to zero.

Solving for *N*(*t*) and *x*(*t*), we get:

From equation (1.2):

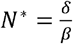 and *x* ^*\**^ = 0 (zero equilibrium point), or 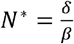 and 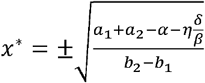 (non-zero equilibrium point).

To study stability and Hopf bifurcation, we derived and analyzed the equation for the properties resulting from the linearization of the system around the equilibrium points. The procedure involved calculating the Jacobian matrix for the delayed system and incorporating the effects of time delays through exponential terms. For equations (1.1) and (1.2), the linearization leads to a transcendental properties equation of the following form:

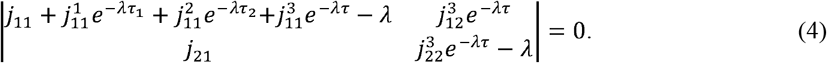

where,

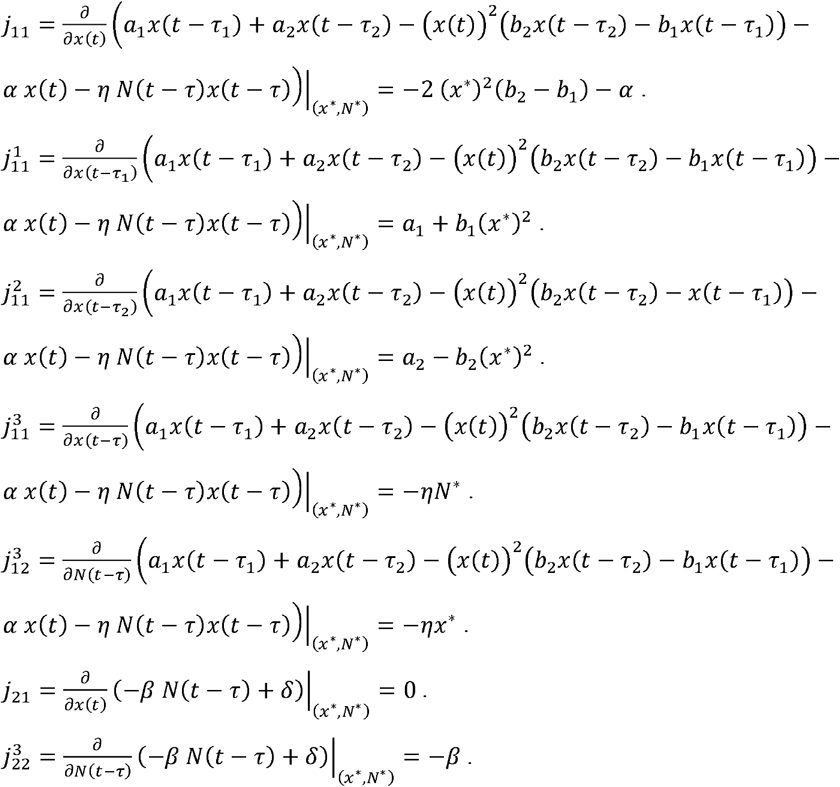

Characteristic equation for zero equilibrium point:

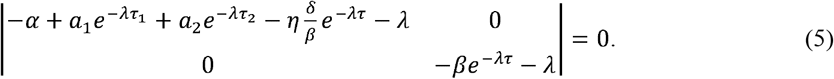

Sinceλ = − *βe* ^-λ τ^, which is always negative, and then the characteristic equation will be:

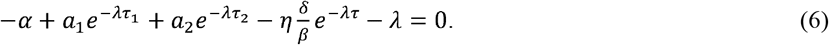

For non-zero equilibrium point:

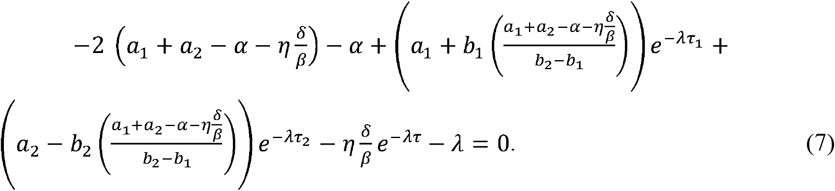

Figure 1 illustrates that α and η play a pivotal role in inducing Hopf bifurcation and keeping stability at the zero-equilibrium point (Fig. 1). Figure 2 demonstrates that the system predominantly experiences Hopf bifurcation across the parameter space at the non-zero equilibrium point during hyperactive oscillations. The dominance of Hopf bifurcation indicates oscillatory behavior driven by parameter interactions (Fig. 2). In figure 3, under the condition α = 0, the system continues to exhibit dominant Hopf bifurcation behavior throughout the parameter space (Fig. 3). The interaction among, *η, δ*, and *β* reveals a consistent pattern of bifurcation, indicating sustained oscillatory dynamics rather than convergence to stability. These results highlight the importance of precisely calibrating system parameters to ensure dynamic stability and robust performance across varying equilibrium conditions.

**Fig. 1:**
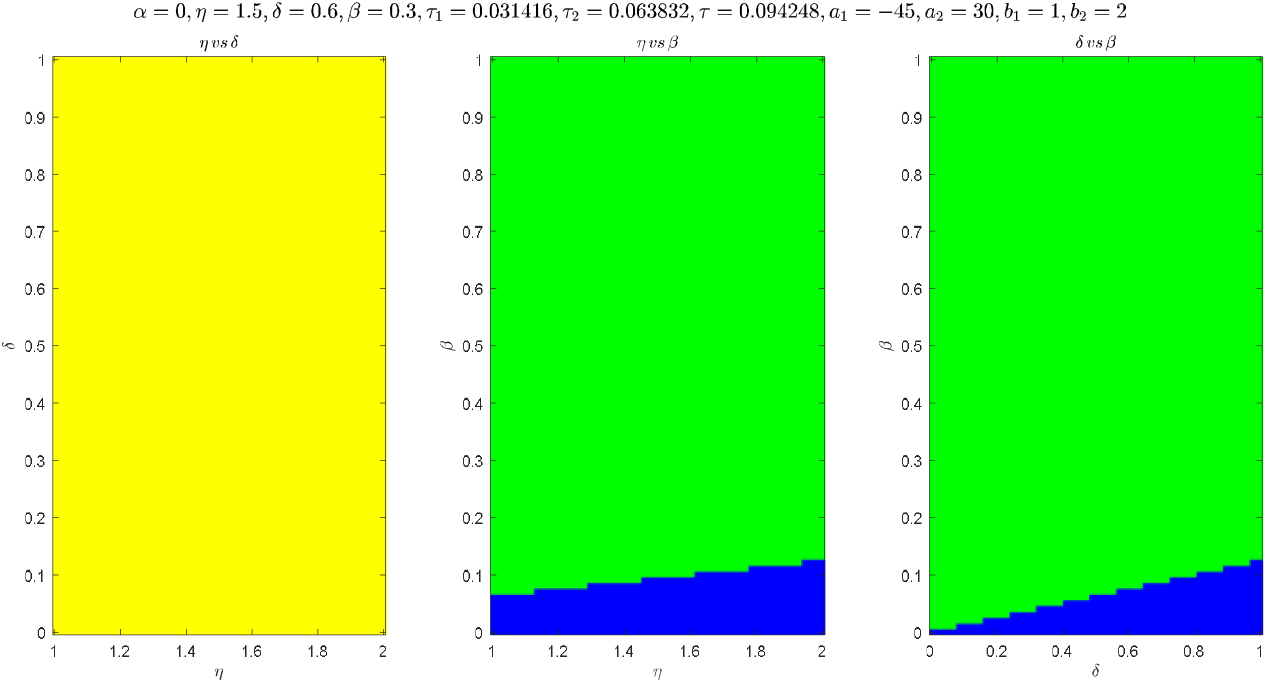
stability (green), instability (blue) and Hopf bifurcation (Yellow) regions between pair parameters for (6).

**Fig. 2:**
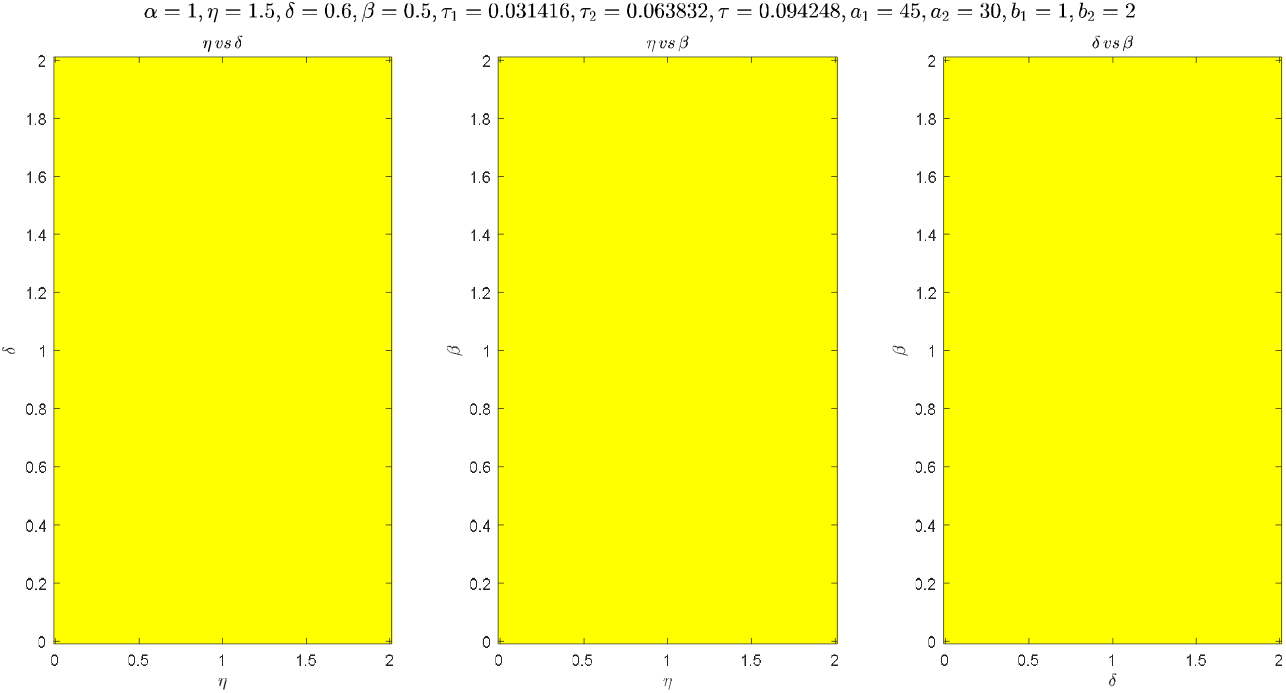
stability (green), instability (blue) and Hopf bifurcation (Yellow) regions between pair parameters for (7).

**Fig. 3:**
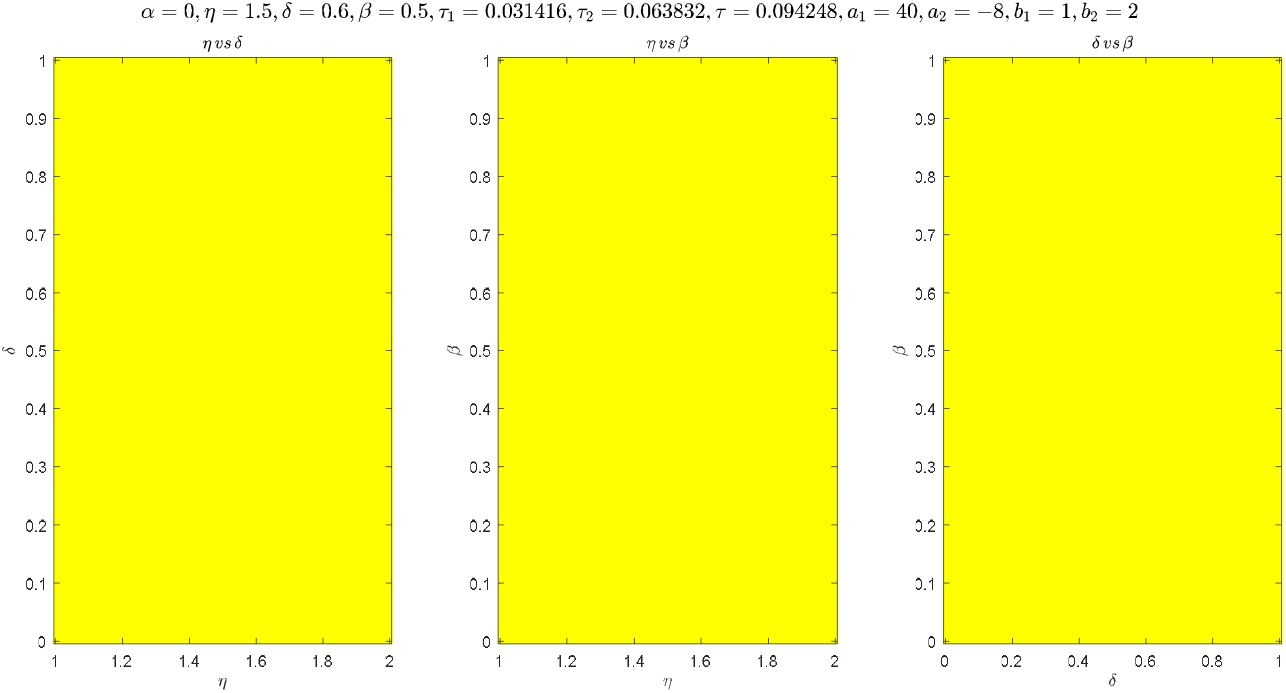
stability (green), instability (blue) and Hopf bifurcation (Yellow) regions between pair parameters for (7).

### Model 2: DBS Effects

The harmonic balance method gives a powerful analytical approach for examining the behavior of the system under deep brain stimulation. To apply this method, we begin by assuming a solution of the form:

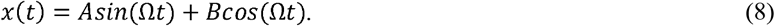

where *A* and *B* are the amplitude coefficients to be determined.

This assumed form captures the fundamental harmonic response of the system to periodic forcing. By putting this expression in (2), we can assess how the system reacts using harmonic parts, which helps us create algebraic equations for *A* and *B*. We then use these equations to analyze the system’s amplitude and frequency behavior under varying stimulation parameters.

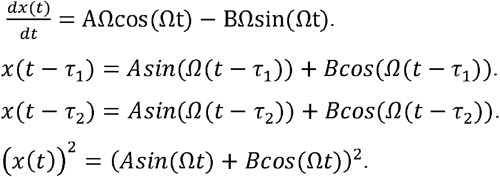

The coefficients of each harmonic (*Cos*(*Ωt*), *sin*(*Ωt*), *Cos*(2*Ωt*), *sin*(2*Ωt*), *Cos*(3*Ωt*) and *sin*(3*Ωt*)) are derived to balance the system under oscillations. These coefficients provide insight into the amplitude, frequency and system dynamics.

### Harmonic Coefficients

For : *Cos*(*Ωt*):

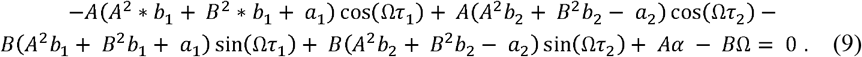

For sin(Ωt):

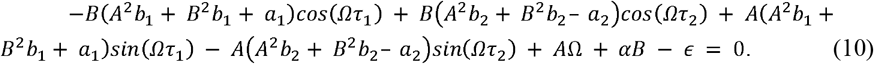

For cos (2Ωt): 0

For sin (2Ωt) :0

For cos (3Ωt) :

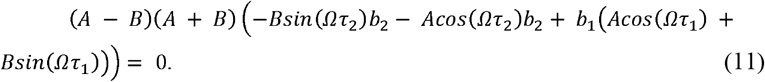

For sin (3Ωt) :

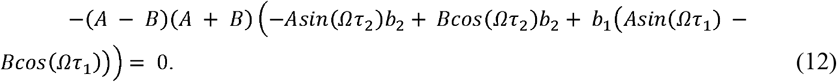

The amplitude modulation,

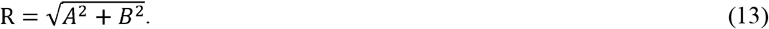

The first harmonic coefficients in (9) and (10) represent the fundamental frequencies in dynamic systems and play an important role in simplifying complex equations. We neglect the higher harmonics, which in many practical cases have a less significant effect. This simplifies the equations and makes the system analysis more clear and solvable. Adding amplitude constraints shows how the system really works without over-amplifying frequencies that aren’t being looked at. This improves the accuracy of the analysis and the reliability of the results and reveals how DBS therapy works.

In the previous three cases, we will replace the parameter values with nanomaterials, as shown in the first model. We will adjust the amplitude to keep it stable and within realistic physical values. Next, we will plot the solutions of |*ϵ* |and *Ω* in the three cases. This procedure helps us understand how the stimulus frequency *Ω* and the amplitude of the external force |*ϵ* | affect the dynamics of the neural system (Fig. 4).

**Fig. 4:**
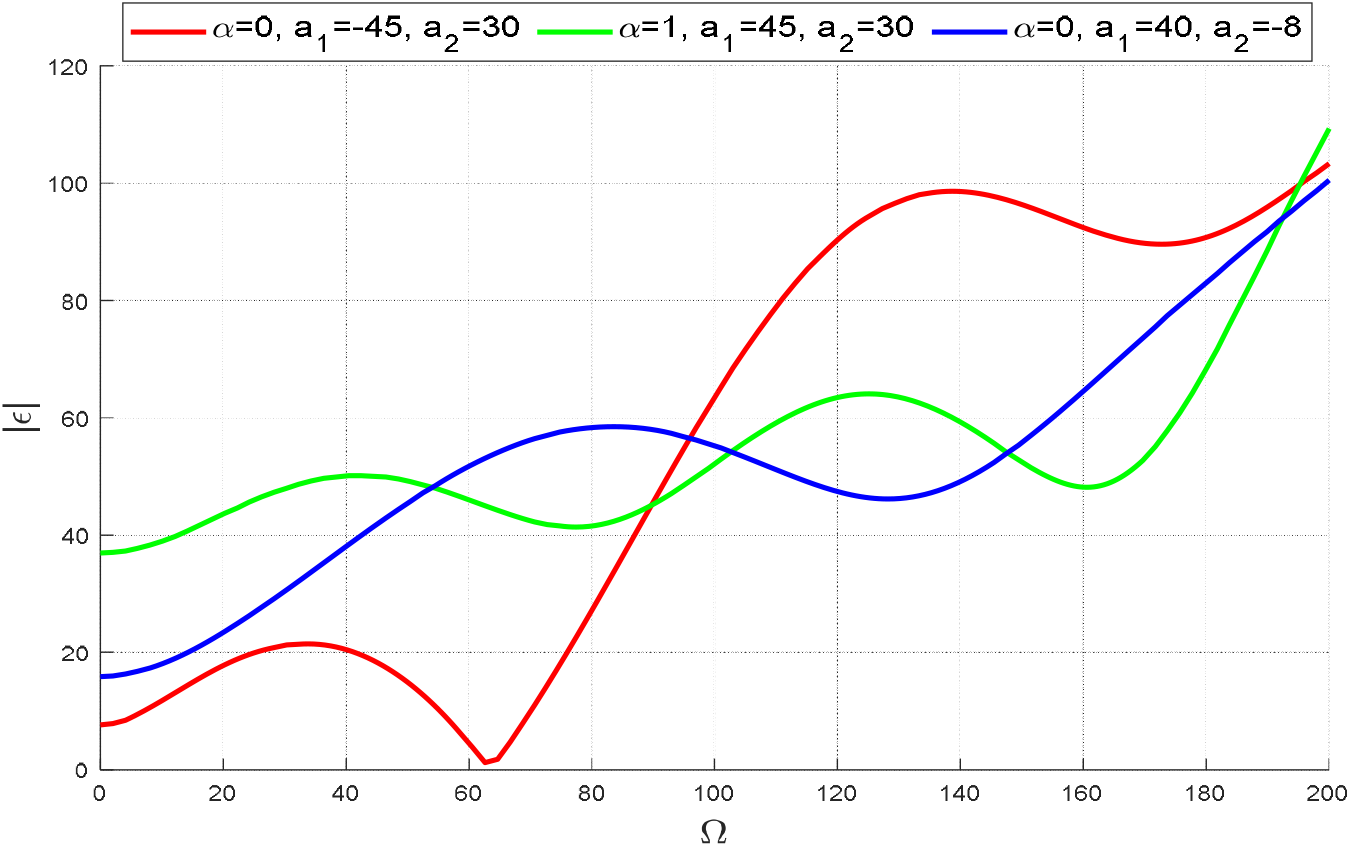
red (*α* = 0, *a*_1_ = − 45 and *a*_2_ = 30), orang (*α* = 0.2, *a*_1_ *=* 45and *a*_2_ = 30) and blue (for α = 0, *a*_1_ = 40 and *a*_1_ = −8) for all 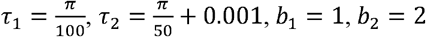.

The symmetry between positive and negative values of e is clearly visible, as using absolute values highlights a balanced system response regardless of the direction of the external force. Symmetry is consistent with figure 5, which analyzes the effects of ϵ and Ω. In figure 5, the effect of different values of ϵ on neural oscillations is highlighted. The results indicate that both positive and negative values of *ϵ* affect frequency and amplitude in the same way, but they influence phase differently, highlighting the need to adjust the synchronization between the various pathways (Fig. 5).

**Fig. 5:**
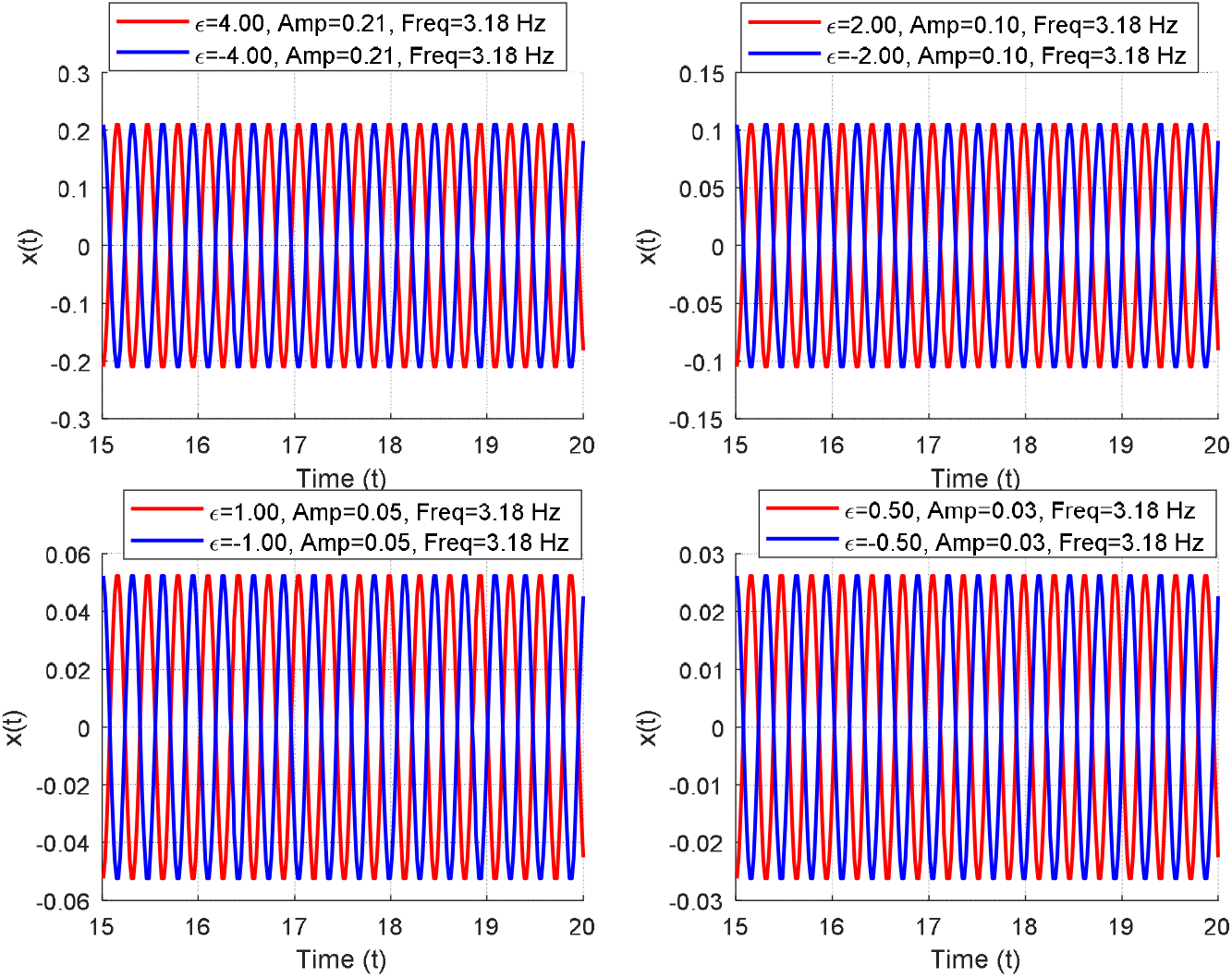
The effect of positive and negative values of *ϵ* on neural oscillations in the first case.

Figure 6 deepens the understanding by examining the effect of different frequencies on neural oscillations, showing the importance of tuning the parameters to achieve the best system response. At low frequencies, neural oscillations are stable but with low amplitude, indicating a limited effect. At intermediate frequencies, significant amplitude amplification is seen, highlighting a strong dynamic match between the stimulating frequency and the natural frequency of the system, which can be interpreted as a partial resonance. ϵ causes amplitude amplification regardless of the direction of the force, highlighting the symmetry that reflects the fundamental nature of the system (Fig. 6).

**Fig. 6:**
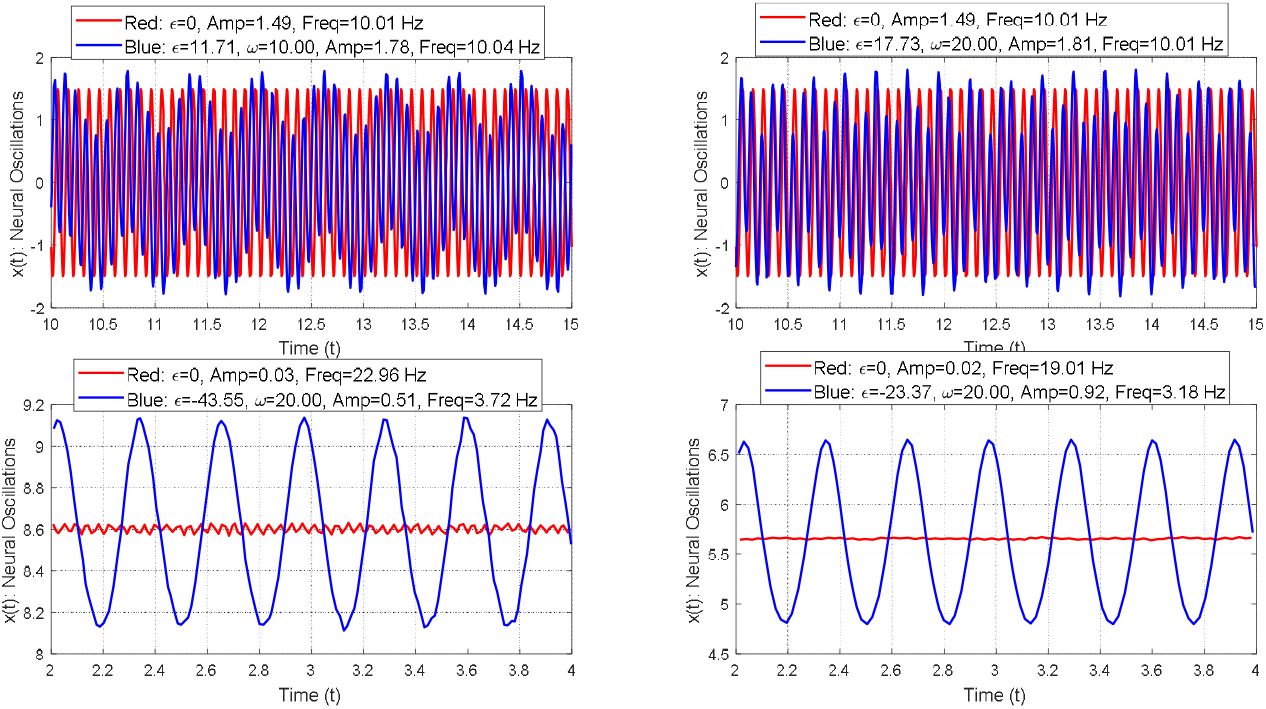
Neural oscillations for different parameter settings. For the first and second subplots: *a*_1_ = − 45, *a*_2_ = 30 and *h*= 4. For the third subplot: *a*_1_ =45, *a*_2_ = 30 and *h*= 4. For the fourth subplot: *a*_1_ =40, *a*_2_ = −8 and *h*= 7.

We can achieve the best balance between minimizing pathological activity and maintaining normal dynamics. Incorrect use of values that are too high or too low can lead to disturbances or weak responses, highlighting the importance of carefully selecting the parameters to ensure the best possible results.

## 4. Numerical Simulation

In this section, we conduct numerical simulations to validate the proposed mathematical models and analyze their behavior under various conditions. The simulations are designed to examine the effects of DBS, nanomaterials and their combined application on pathological oscillations within the basal ganglia. We employ default parameter values to represent the intrinsic dynamics of the neural pathways and the pathological consequences associated with increased response delays in these circuits.

### Model 1: Nanomateriasl Effects

Figure 7 illustrates the effect of varying parameters (*a*_1_ = −45, *a*_2_ = 30, *b*_1_ =1,*b*_2_ =2 and *α* = 0) on neural oscillations within the basal ganglia pathways. The results indicate that a low value of *β* = 0.1, combined with higher values of *η* = 2.0 and *δ*=1.0, leads to a significant reduction in oscillation amplitude while maintaining a frequency of approximately 10.86 Hz. This finding demonstrates the stabilizing effect of nanomaterials on the system. Conversely, when *β* = 0.6 is increased and *η* = 0.5, δ =0.4 are reduced, a modest decrease in oscillation amplitude is observed, with a maintained frequency of 10.04 *Hz*. This pattern reflects the influence of nanomaterials on amplitude modulation without disrupting the oscillatory frequency (Fig. 7). Figure 8 explores the effect of the nanomaterials parameters *β, η*, and *δ* in a second case, where motor cortex activity is increased by modifying the values to *a*_1_= 45, *a*_2_ = 30 and *α*= 1. The results are consistent with those in figure 7: although the amplitude remains stable, motor cortex activity is reduced while frequency remains nearly constant (Fig. 8). This highlights the nanomaterials amplitude to stabilize the system without significantly disrupting its natural oscillatory dynamics. Figure 9 examines a third case involving further adjustments to the core parameters, specifically *a*_1_ = 40, *a*_2_ =−8, and *α* = 0. The results match those in figures 7 and 8, confirming that nanomaterials consistently help maintain natural oscillation and stability, even with different parameter settings (Fig. 9).

**Fig. 7:**
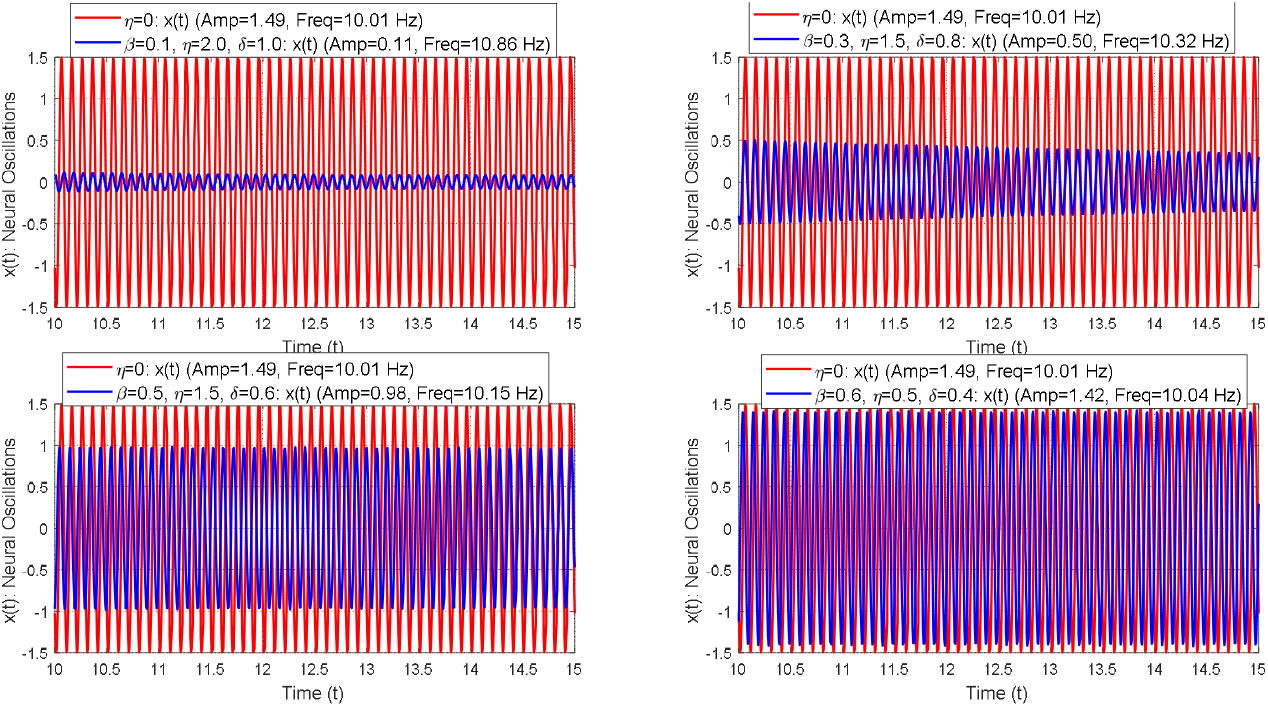
Neural oscillations under varying parameters for the basal ganglia pathways. The base parameters (*a*_1_ = −45, *a*_2_ = 30, *b*_1_ = 1, *b*_2_ = 2 and *α* = 0).

**Fig. 8:**
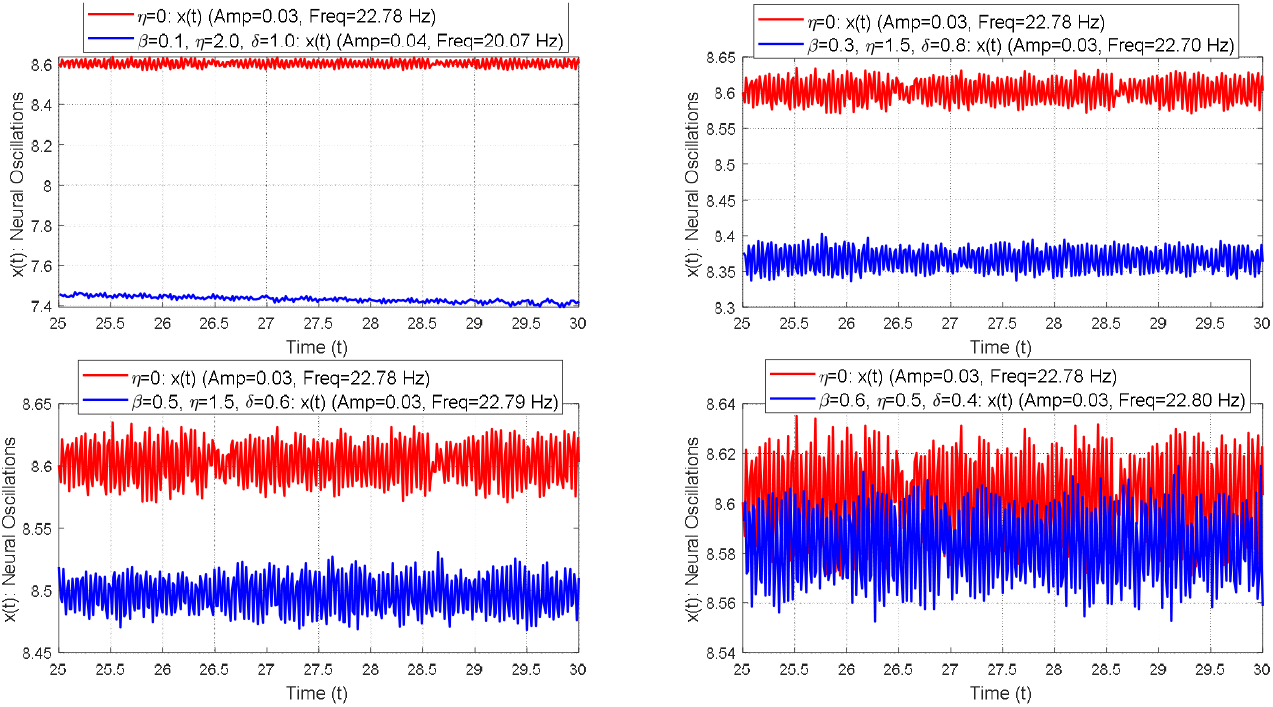
Neural oscillations under varying parameters for the basal ganglia pathways. The base parameters (*a*_1_ = 45, *a*_2_ = 30, *b*_1_ = 1, *b*_2_ = 2 and *α* = 1).

**Fig. 9:**
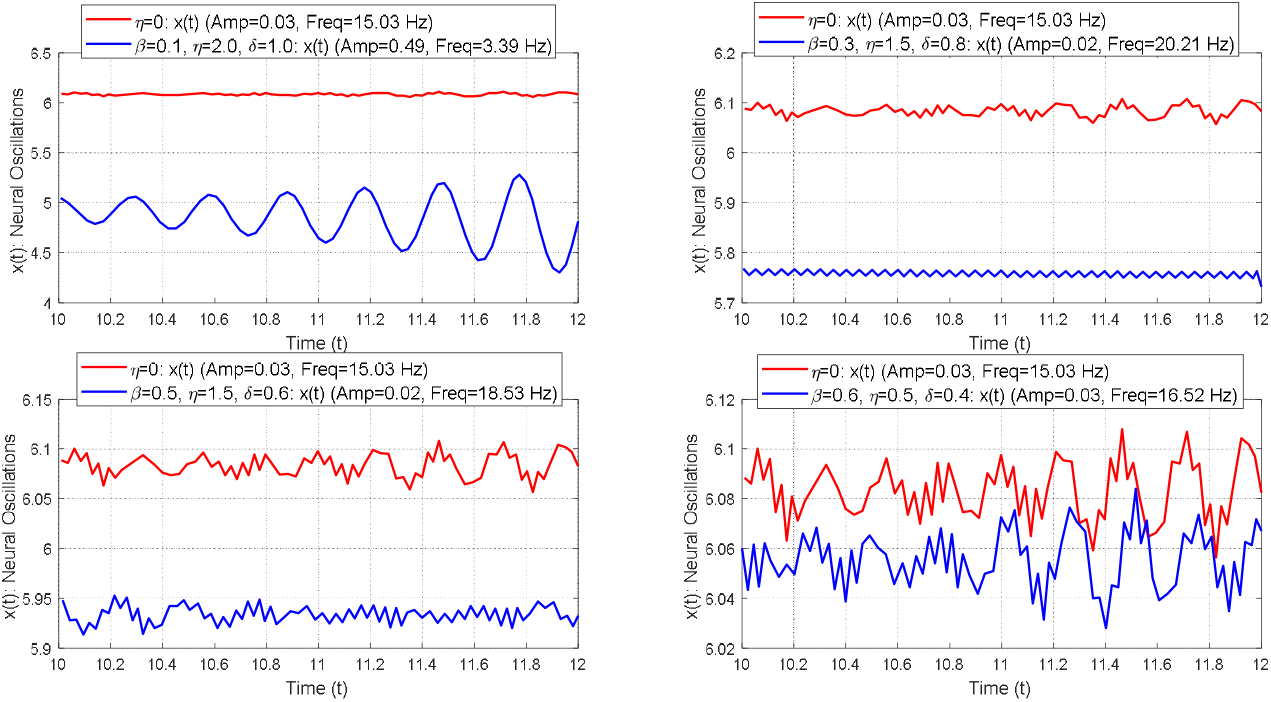
Neural oscillations under varying parameters for the basal ganglia pathways. The base parameters (*a*_1_ = 40, *a*_2_ =−8, *b*_1_ = 1, *b*_2_ = 2 and *α* = 0).

### Model 2: DBS Effects

Figure 10 examines the effect of varying *ϵ* under fixed stimulation frequency (*Ω*=20 *Hz*) on neural oscillations during DBS. The key parameters are set to *a*_1_ = −45, *a*_2_= 30, *b*_1_ = 1, *b*_2_ = 2 and *α* = 0. As *ϵ* increases (10, 20, 30, and 40), the oscillation amplitude shows a slight initial rise, indicating a dynamic system response. However, at *ϵ* = 40, the frequency drops significantly to 3.16 *Hz*, revealing the system’s tendency toward overdamping under excessive stimulation. These findings underscore the importance of fine-tuning *ϵ* to suppress pathological oscillations effectively while preserving natural frequency behavior (Fig. 10). Figure 11 explores the combined effect of *ϵ* and *Ω* on neural oscillations under DBS. While the base parameters are retained, modifications are applied: *a*_1_= 45, *a*_2_ = 30, *α* = 1, *ϵ* = {0.2,0.5,1,4} and *Ω* = 20 *Hz*. Across these settings, a marked reduction in frequency is observed, accompanied by the suppression of pathological beta oscillations. However, this comes with a notable increase in oscillation amplitude. These results illustrate the delicate trade-off in DBS optimization—excessive stimulation may inadvertently suppress normal neural activity, potentially affecting motor function (Fig. 11). Figure 12 further evaluates the influence of *ϵ* and *Ω* on oscillatory dynamics under new parameter settings: *a*_1_ = 40, *a2=* −8, *α* = 0 and *ϵ* ={0,1.0.3,1,2} The outcomes are consistent with those in figure 11, frequency suppression is accompanied by increased amplitude. These results highlight the importance of carefully adjusting *ϵ* to gain therapeutic benefits while keeping the natural frequency patterns of basal ganglia oscillations intact (Fig. 12).

**Fig. 10:**
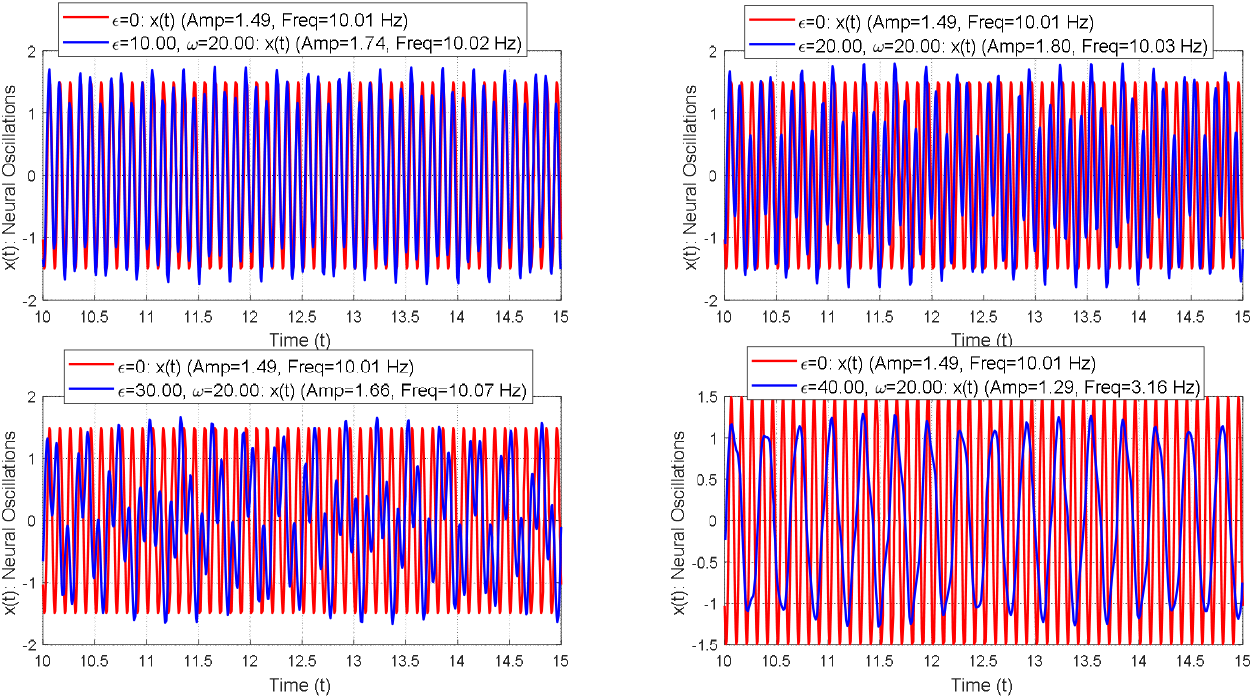
Neural oscillations under varying parameters for the basal ganglia pathways. The base parameters (*a*_1_*=* −45, *a*_2_ = 30, *b*_1_ = 1, *b*_2_ = 2 and *α* = 0).

**Fig. 11:**
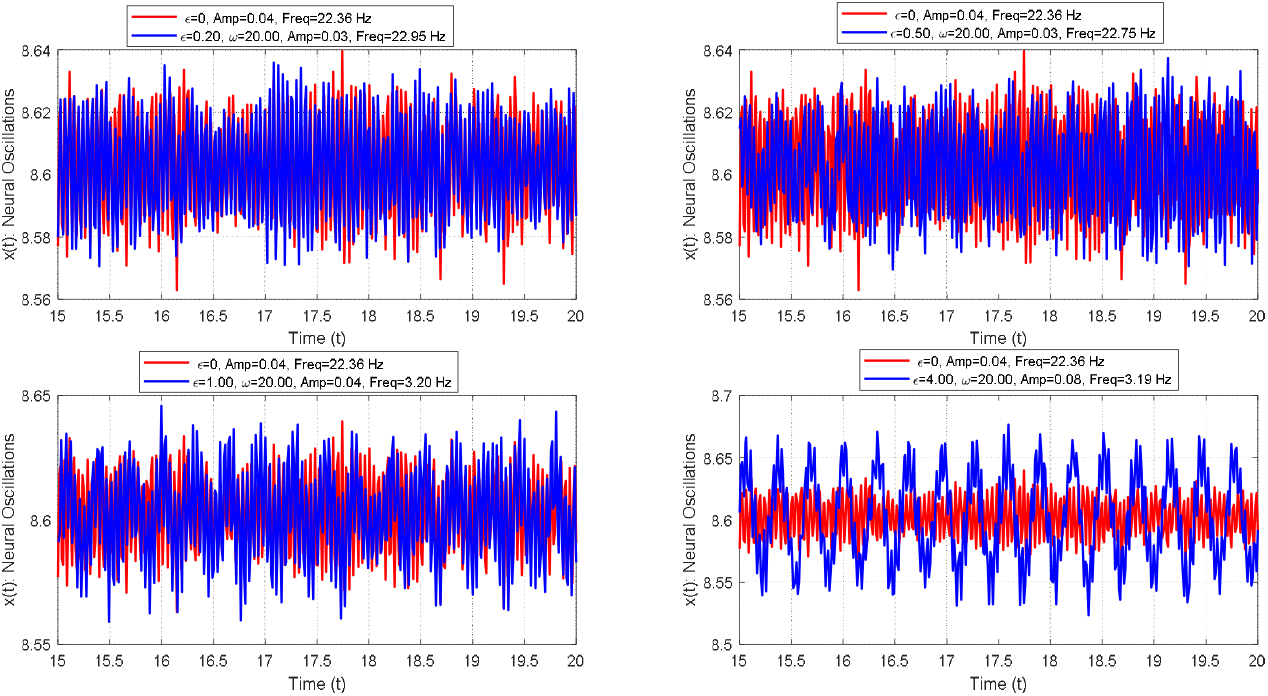
Neural oscillations under varying parameters for the basal ganglia pathways. The base parameters (*a*_1_*=* 45, *a*_2_ = 30, *b*_1_ = 1, *b*_2_ = 2 and *α* = 0).

**Fig. 12:**
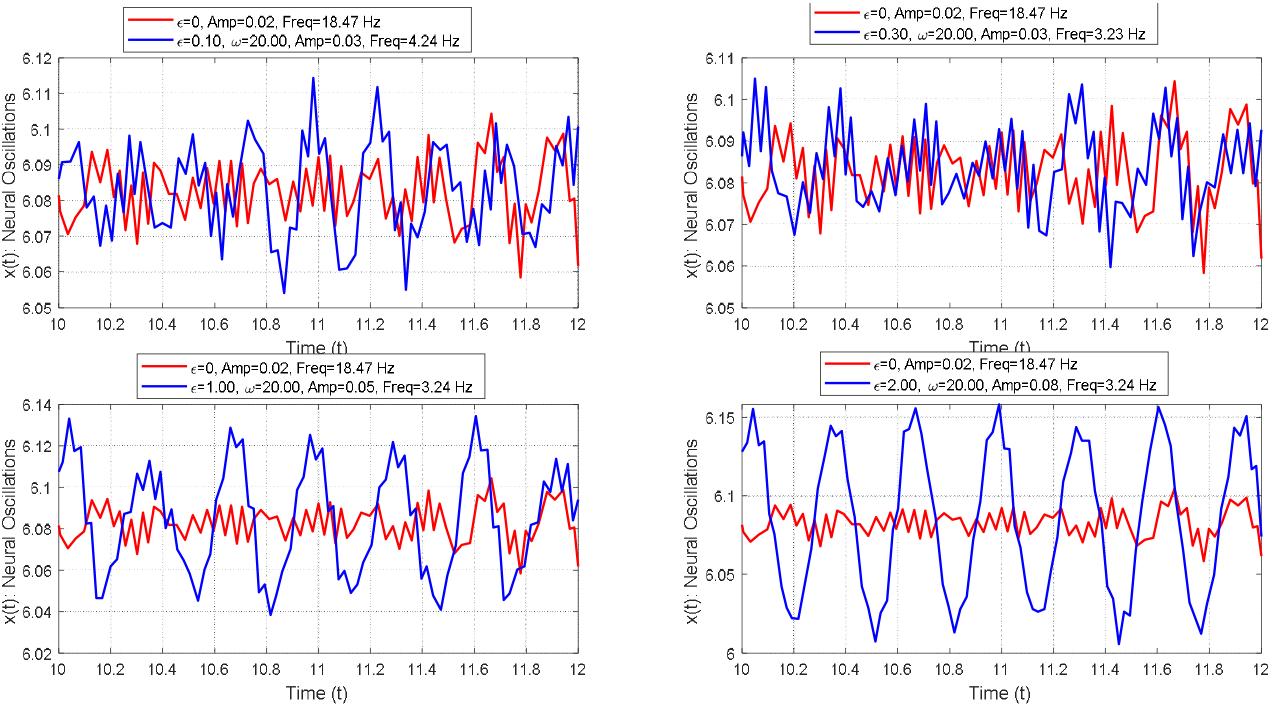
Neural oscillations under varying parameters for the basal ganglia pathways. The base parameters (*a*_1_*=* 40, *a*_2_ = −8, *b*_1_ = 1, *b*_2_ = 2 and *α* = 0).

### Model 3: Combined Effects of Nanomaterials and DBS

We integrated the effect of nanomaterials into the DBS models to explore strategies for enhancing their performance and improving the stability of neural dynamics. The outcomes were compared to the previously obtained results in figures 10, 11, and 12 to evaluate the added benefits of combining interventions, as illustrated in figures 13, 14, and 15. Figure 13 demonstrates the influence of nanomaterials on neural oscillatory behavior. In contrast to figure 10, incorporating nanomaterials led to improved consistency and a notable reduction in oscillation amplitude at higher values of *ϵ*, suggesting enhanced control over overstimulation effects. Figure 14, when compared to figure 11, shows a substantial decrease in motor cortex hyperactivity while maintaining low-frequency oscillations, indicating a therapeutic gain without compromising oscillatory integrity. Similarly, figure 15 reveals a reduction in motor cortex activity relative to figure 12, reaffirming the stabilizing effect of nanomaterials on both amplitude and frequency dynamics. Overall, integrating nanomaterials into DBS frameworks enhances the stability of neural oscillations and effectively reduces pathological activity, thereby contributing to improved therapeutic outcomes.

**Fig. 13:**
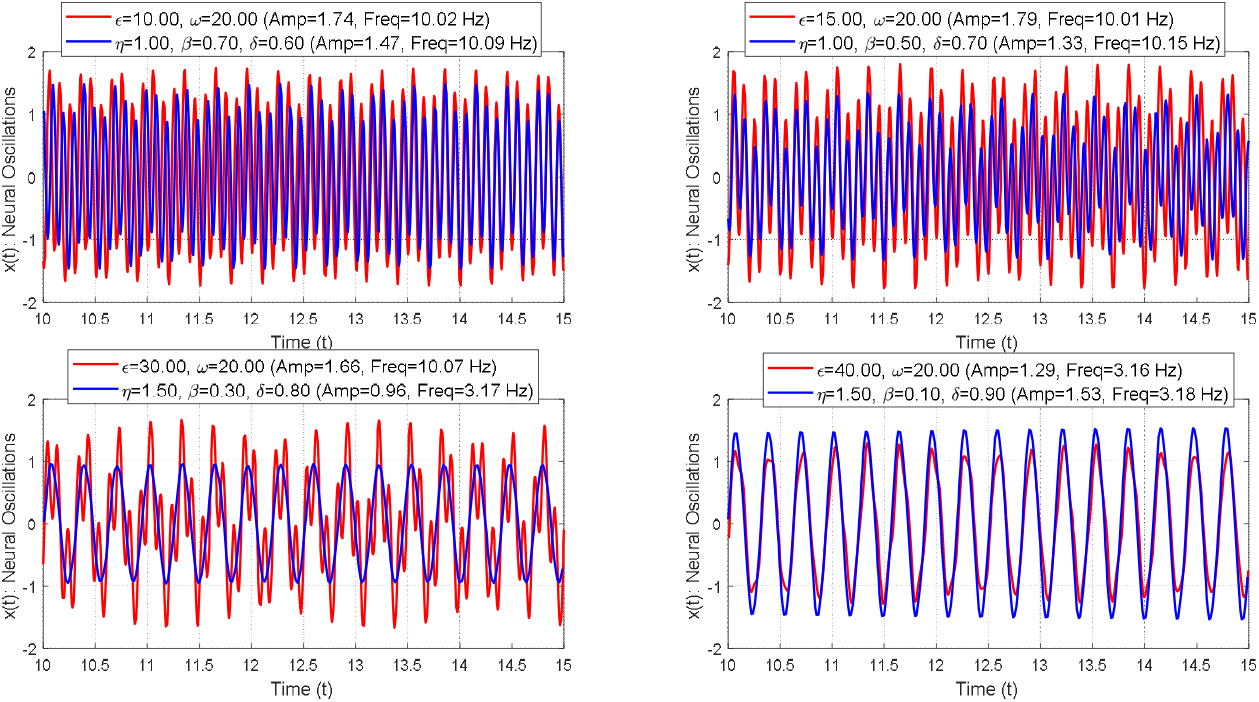
Neural oscillations under varying parameters for the basal ganglia pathways. The base parameters (*a*_1_*=* −45, *a*_2_ = 30, *b*_1_ = 1, *b*_2_ = 2 and *α* = 0).

**Fig. 14:**
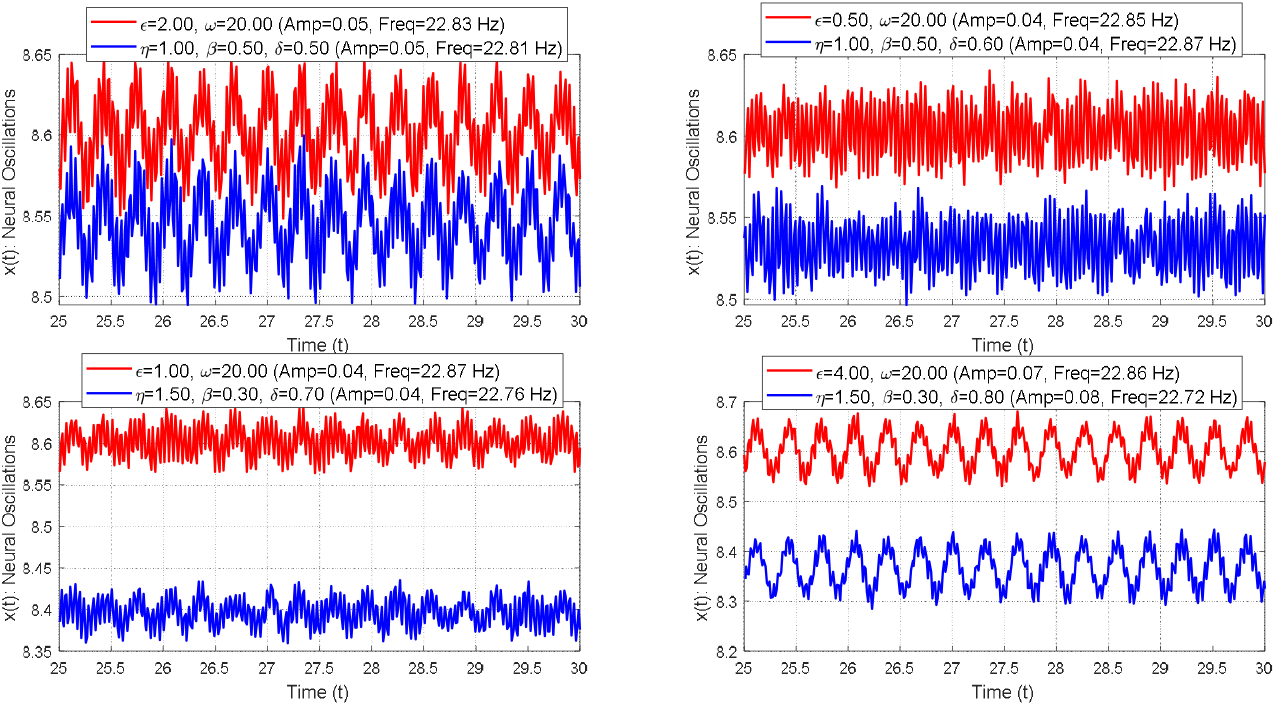
Neural oscillations under varying parameters for the basal ganglia pathways. The base parameters (*a*_1_*=* 45, *a*_2_ = 30, *b*_1_ = 1, *b*_2_ = 2 and *α* = 1).

**Fig. 15:**
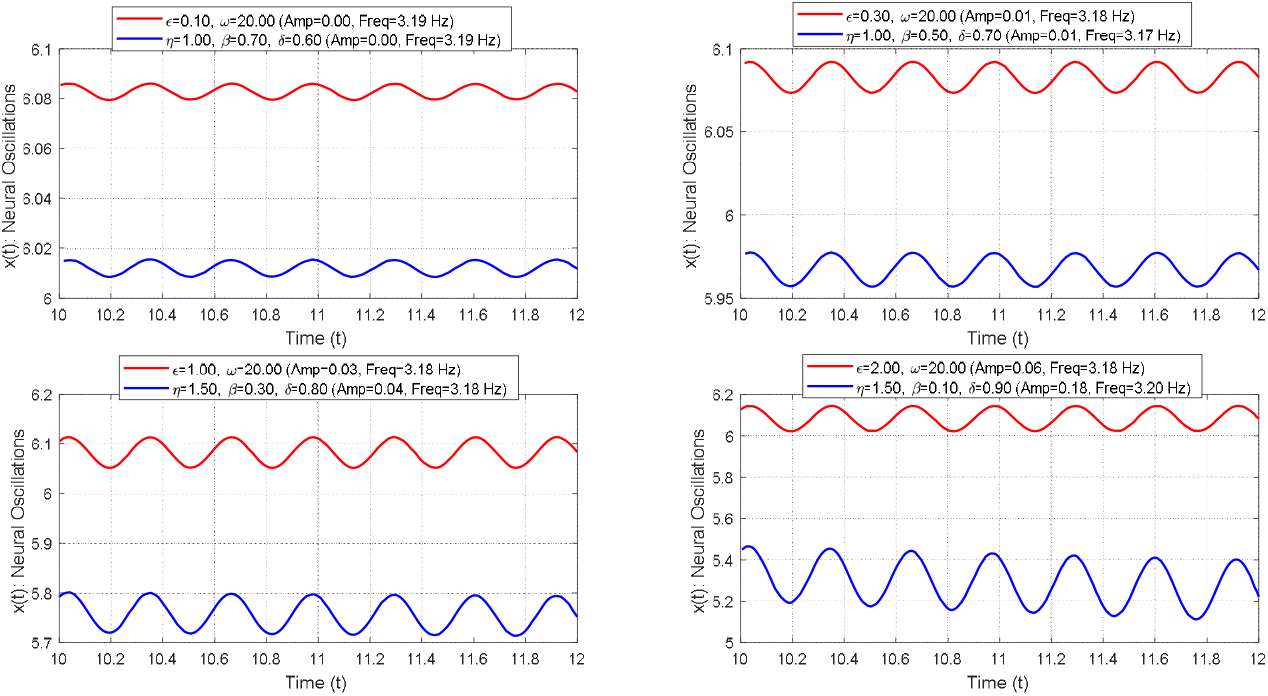
Neural oscillations under varying parameters for the basal ganglia pathways. The base parameters (*a*_1_*=* 40, *a*_2_ = −8, *b*_1_ = 1, *b*_2_ = 2 and *α* = 0).

## 6. Discussion

The DBS-only models presented in figures 10, 11, and 12 provide critical insights into how stimulation affects neural oscillation dynamics. When *ϵ* is low, both amplitude and frequency remain relatively unchanged, reflecting a typical neural response. However, as *ϵ* increases, particularly at *ϵ* = 40, a marked reduction in frequency to 3.16 Hz is observed, indicating an overdamping effect due to excessive stimulation. While DBS can effectively suppress pathological beta-band oscillations, the accompanying increase in amplitude may lead to hyperactivity within the motor cortex.

In contrast, the incorporation of nanomaterials into the DBS framework, as shown in figures 13, 14, and 15, enhances the stability of neural dynamics. The combination reduces pathological amplitude and improves frequency stability, particularly at high values of *ϵ*. Nanomaterials also help mitigate motor cortex hyperactivity while preserving the brain’s natural oscillatory frequencies.

The integration of intrinsic and extrinsic frequencies allows for finer control over neural dynamics, contributing to improved therapeutic performance. In the DBS-only models, partial resonances were observed at certain frequencies (*Ω* = 20 *Hz*), where periodic stimulation amplified pathological amplitudes. Although such resonance may help suppress pathological frequencies, it can also destabilize the system under certain conditions.

The addition of nanomaterials acts as a damping mechanism, reducing amplitude amplification and thus mitigating the risks associated with resonance-induced instability. This improved dynamic balance fosters more stable neural oscillations and enhances therapeutic outcomes by simultaneously reducing pathological activity and maintaining physiological signal integrity.

## 6. Conclusion

This study offers a scientific way to understand and improve brain wave patterns by combining nanomaterials with deep brain stimulation. The findings highlight both the potential and the limitations of this combined approach. While the deep brain stimulation-only model effectively attenuates pathological frequencies, excessive stimulation leads to over-damping, evidenced by a significant frequency reduction and may destabilize motor cortex activity due to amplitude amplification. These results underscore the importance of careful parameter tuning.

Incorporating nanomaterials into the system significantly improves neural stability. Their damping effect reduces pathological amplitudes while preserving physiological frequency ranges. This two-part approach lowers harmful vibrations while keeping normal movements, helping to calm the motor cortex, improve movement control, and lessen possible side effects. The results demonstrate that this hybrid strategy can balance pathological suppression with the preservation of normal neural function.

However, excessive use of nanomaterials could suppress normal neural activity, potentially impairing fine motor control. Since this study is based on mathematical modeling and numerical simulations, experimental validation is essential to confirm the findings and assess their clinical applicability.

Combining nanomaterials with deep brain stimulation offers a new and hopeful way to treat conditions like Parkinson’s disease that involve brain wave problems. This research is a significant move towards creating tailored neuromodulation methods that consider each person’s brain activity, and it shows the importance of future studies in living organisms to investigate how effective and safe these methods are over time.

## Funding

No funding exists.

## Ethical Approval

There is no clinical data. human or animal experiments. The manuscript’s data were suggested by the author to explain the model’s behavior and the generation of neural impulses.

## Availability of data and materials

The datasets generated and/or analyzed during the current study are available from the corresponding author upon reasonable request.

## Notes

### Competing Interest Statement

The authors have declared no competing interest.

## References

[1] Rocha, G. S., Freire, M. a. M., Britto, A. M., Paiva, K. M., Oliveira, R. F., Fonseca, I. a. T., Araújo, D. P., Oliveira, L. C., Guzen, F. P., Morais, P. L. a. G., & Cavalcanti, J. R. L. P. (2023). Basal ganglia for beginners: the basic concepts you need to know and their role in movement control. Frontiers in Systems Neuroscience, 17. 10.3389/fnsys.2023.1242929

[2] Ikemoto, S., Yang, C., & Tan, A. (2015). Basal ganglia circuit loops, dopamine and motivation: A review and enquiry. Behavioural Brain Research, 290, 17–31. 10.1016/j.bbr.2015.04.018

[3] Graybiel, A. M. (2008). Habits, rituals, and the evaluative brain. Annual Review of Neuroscience, 31(1), 359–387. 10.1146/annurev.neuro.29.051605.112851

[4] De Virgilio, A., Greco, A., Fabbrini, G., Inghilleri, M., Rizzo, M. I., Gallo, A., Conte, M., Rosato, C., Appiani, M. C., & De Vincentiis, M. (2016). Parkinson’s disease: Autoimmunity and neuroinflammation. Autoimmunity Reviews, 15(10), 1005–1011. 10.1016/j.autrev.2016.07.022

[5] Kouli, A., Torsney, K. M., & Kuan, W. (2018). Parkinson’s Disease: etiology, neuropathology, and pathogenesis. In Codon Publications eBooks (pp. 3–26). 10.15586/codonpublications.parkinsonsdisease.2018.ch1

[6] Fraser, H., & Brown, C. G. (2014). Britain since 1707. In Routledge eBooks. 10.4324/9781315835310

[7] Hariz, M., & Blomstedt, P. (2022). Deep brain stimulation for Parkinson’s disease. Journal of Internal Medicine, 292(5), 764–778. 10.1111/joim.13541

[8] De Oliveira, F., Vaz, R., Chamadoira, C., Rosas, M. J., & Ferreira-Pinto, M. J. (2023). Bilateral deep brain stimulation of the subthalamic nucleus: Targeting differences between the first and second side. Neurocirugía (English Edition), 34(4), 186–193. 10.1016/j.neucie.2022.07.001

[9] Lozano, A. M., Lipsman, N., Bergman, H., Brown, P., Chabardes, S., Chang, J. W., Matthews, K., McIntyre, C. C., Schlaepfer, T. E., Schulder, M., Temel, Y., Volkmann, J., & Krauss, J. K. (2019). Deep brain stimulation: current challenges and future directions. Nature Reviews Neurology, 15(3), 148–160. 10.1038/s41582-018-0128-2

[10] Eldaief, M. C., Dickerson, B. C., & Camprodon, J. A. (2022). Transcranial magnetic stimulation for the neurological patient: scientific principles and applications. Seminars in Neurology, 42(02), 149–157. 10.1055/s-0041-1742265

[11] Kühn, A. A., & Volkmann, J. (2016). Innovations in deep brain stimulation methodology. Movement Disorders, 32(1), 11–19. 10.1002/mds.26703

[12] Zarzycki, M. Z., & Domitrz, I. (2019). Stimulation-induced side effects after deep brain stimulation – a systematic review. Acta Neuropsychiatrica, 32(2), 57–64. 10.1017/neu.2019.35

[13] Yang, S., Baeg, E., Kim, K., Kim, D., Xu, D., Ahn, J., & Yang, S. (2023). Neurodiagnostic and neurotherapeutic potential of graphene nanomaterials. Biosensors and Bioelectronics, 247, 115906. 10.1016/j.bios.2023.115906

[14] Malik, J. A., AjgarAnsari, J., Ahmed, S., Rani, A., Ansari, S. Y., & Anwar, S. (2023). Emerging selenium nanoparticles for CNS intervention. In Biomedical engineering. 10.5772/intechopen.109418

[15] Zecchi, S., Cristoforo, G., Piatti, E., Torsello, D., Ghigo, G., Tagliaferro, A., Rosso, C., & Bartoli, M. (2024). A concise review of recent advancements in carbon nanotubes for aerospace applications. Micromachines, 16(1), 53. 10.3390/mi16010053

[16] Yuwen, T., Shu, D., Zou, H., Yang, X., Wang, S., Zhang, S., Liu, Q., Wang, X., Wang, G., Zhang, Y., & Zang, G. (2023). Carbon nanotubes: a powerful bridge for conductivity and flexibility in electrochemical glucose sensors. Journal of Nanobiotechnology, 21(1). 10.1186/s12951-023-02088-7

[17] Seyedkhani, S. A. (2024). Carbon nanomaterials for neural interfaces: synthesis, properties and applications. In IntechOpen eBooks. 10.5772/intechopen.1006603

[18] Shabani, L., Abbasi, M., Azarnew, Z., Amani, A. M., & Vaez, A. (2023). Neuro-nanotechnology: diagnostic and therapeutic nano-based strategies in applied neuroscience. BioMedical Engineering OnLine, 22(1). 10.1186/s12938-022-01062-y

[19] Riva, E. R., Özkan, M., Stellacci, F., & Micera, S. (2024). Combining external physical stimuli and nanostructured materials for upregulating pro-regenerative cellular pathways in peripheral nerve repair. Frontiers in Cell and Developmental Biology, 12. 10.3389/fcell.2024.1491260

[20] Kulkarni, M., Patel, K., Patel, A., Patel, S., Desai, J., Patel, M., Shah, U., Patel, A., & Solanki, N. (2023). Nanomaterials as drug delivery agents for overcoming the blood-brain barrier: A comprehensive review. ADMET & DMPK. 10.5599/admet.2043

[21] Liu, J., Wang, T., Dong, J., & Lu, Y. (2025). The blood–brain barriers: novel nanocarriers for central nervous system diseases. Journal of Nanobiotechnology, 23(1). 10.1186/s12951-025-03247-8

[22] Razavi, Z. S., Alizadeh, S. S., Razavi, F. S., Souri, M., & Soltani, M. (2025). Advancing neurological disorders therapies: Organic nanoparticles as a key to blood-brain barrier penetration. International Journal of Pharmaceutics, 670, 125186. 10.1016/j.ijpharm.2025.125186

[23] Song, Q., Li, J., Li, T., & Li, H. (2024). Nanomaterials that Aid in the Diagnosis and Treatment of Alzheimer’s Disease, Resolving Blood–Brain Barrier Crossing Ability. Advanced Science, 11(38). 10.1002/advs.202403473

[24] Goma, A. A., Salama, A. R., Tohamy, H. G., Rashed, R. R., Shukry, M., & El-Kazaz, S. E. (2024). Examining the Influence of Zinc Oxide Nanoparticles and Bulk Zinc Oxide on Rat Brain Functions: a Comprehensive Neurobehavioral, Antioxidant, Gene Expression, and Histopathological Investigation. Biological Trace Element Research, 202(10), 4654–4673. 10.1007/s12011-023-04043-x

[25] Mohammad, N., Khan, U. A., Warsi, M. H., Alkreathy, H. M., Karim, S., Jain, G. K., & Ali, A. (2023). Intranasal cerium oxide nanoparticles improves locomotor activity and reduces oxidative stress and neuroinflammation in haloperidol-induced parkinsonism in rats. Frontiers in Pharmacology, 14. 10.3389/fphar.2023.1188470

[26] Hernando, S., SantoslVizcaíno, E., Igartua, M., & Hernandez, R. M. (2023). Targeting the central nervous system: From synthetic nanoparticles to extracellular vesicles—Focus on Alzheimer’s and Parkinson’s disease. Wiley Interdisciplinary Reviews Nanomedicine and Nanobiotechnology, 15(5). 10.1002/wnan.1898

[27] Ahmed-Farid, O. A., Taha, M., Bakeer, R. M., Radwan, O. K., Hendawy, H. a. M., Soliman, A. S., & Yousef, E. (2021). Effects of bee venom and dopamine-loaded nanoparticles on reserpine-induced Parkinson’s disease rat model. Scientific Reports, 11(1). 10.1038/s41598-021-00764-y

[28] Kheiriabad, S., Jafari, A., Aghdash, S. N., Dolatabadi, J. E. N., Andishmand, H., & Jafari, S. M. (2024). Applications of advanced nanomaterials in biomedicine, pharmaceuticals, agriculture, and food industry. BioNanoScience, 14(4), 4298–4321. 10.1007/s12668-024-01506-w

[29] Alahi, M. E. E., Rizu, M. I., Tina, F. W., Huang, Z., Nag, A., & Afsarimanesh, N. (2023). Recent advancements in Graphene-Based implantable electrodes for neural Recording/Stimulation. Sensors, 23(24), 9911. 10.3390/s23249911

[30] Ramezani, M., Kim, J., Liu, X., Ren, C., Alothman, A., De-Eknamkul, C., Wilson, M. N., Cubukcu, E., Gilja, V., Komiyama, T., & Kuzum, D. (2024). High-density transparent graphene arrays for predicting cellular calcium activity at depth from surface potential recordings. Nature Nanotechnology, 19(4), 504–513. 10.1038/s41565-023-01576-z

[31] Wang, Y., Yang, X., Zhang, X., Wang, Y., & Pei, W. (2023). Implantable intracortical microelectrodes: reviewing the present with a focus on the future. Microsystems & Nanoengineering, 9(1). 10.1038/s41378-022-00451-6

[32] Wang, T., Chen, Y., Wang, Y., Lee, S., Lee, Y., & Dong, J. (2024). Advanced Neural Probe Sensors toward MultilModal Sensing and Modulation: Design, Integration, and Applications. Advanced Sensor Research. 10.1002/adsr.202400142

[33] Gou, S., Yang, S., Cheng, Y., Yang, S., Liu, H., Li, P., & Du, Z. (2024). Applications of 2D nanomaterials in neural interface. International Journal of Molecular Sciences, 25(16), 8615. 10.3390/ijms25168615

[34] Viana, D., Walston, S. T., Masvidal-Codina, E., Illa, X., Rodríguez-Meana, B., Del Valle, J., Hayward, A., Dodd, A., Loret, T., Prats-Alfonso, E., De La Oliva, N., Palma, M., Del Corro, E., Del Pilar Bernicola, M., Rodríguez-Lucas, E., Gener, T., De La Cruz, J. M., Torres-Miranda, M., Duvan, F. T., … Garrido, J. A. (2024). Nanoporous graphene-based thin-film microelectrodes for in vivo high-resolution neural recording and stimulation. Nature Nanotechnology, 19(4), 514–523. 10.1038/s41565-023-01570-5

[35] Krishnan, S. K., Nataraj, N., Meyyappan, M., & Pal, U. (2023). Graphene-Based Field-Effect Transistors in biosensing and Neural interfacing Applications: Recent Advances and Prospects. Analytical Chemistry, 95(5), 2590–2622. 10.1021/acs.analchem.2c03399

[36] Liu, X., Gong, Y., Jiang, Z., Stevens, T., & Li, W. (2024). Flexible high-density microelectrode arrays for closed-loop brain–machine interfaces: a review. Frontiers in Neuroscience, 18. 10.3389/fnins.2024.1348434

[37] Wang, X., Wang, S., Lu, Y., Wang, Y., Song, Y., Wang, X., & Nyamgerelt, M. (2022). Graphene and graphene-based materials in axonal repair of spinal cord injury. Neural Regeneration Research, 17(10), 2117. 10.4103/1673-5374.335822

[38] Villa, J., Cury, J., Kessler, L., Tan, X., & Richter, C. (2024). Enhancing biocompatibility of the brain-machine interface: A review. Bioactive Materials, 42, 531–549. 10.1016/j.bioactmat.2024.08.034

[39] Xie, Y., Peng, Y., Guo, J., Liu, M., Zhang, B., Yin, L., Ding, H., & Sheng, X. (2024). Materials and devices for high-density, high-throughput micro-electrocorticography arrays. Fundamental Research, 5(1), 17–28. 10.1016/j.fmre.2024.01.016

[40] Liu, X., Gong, Y., Jiang, Z., Stevens, T., & Li, W. (2024b). Flexible high-density microelectrode arrays for closed-loop brain–machine interfaces: a review. Frontiers in Neuroscience, 18. 10.3389/fnins.2024.1348434

[41] Zhang, H., Rong, G., Bian, S., & Sawan, M. (2022). Lab-on-Chip Microsystems for Ex vivo Network of Neurons Studies: A review. Frontiers in Bioengineering and Biotechnology, 10. 10.3389/fbioe.2022.841389

[42] Agiza, H. N., Sohaly, M. A., & Elfouly, M. A. (2022). Small two-delay differential equations for Parkinson’s disease models using Taylor series transform. Indian Journal of Physics, 97(1), 39–46. 10.1007/s12648-021-02263-2

[43] Park, J., Yoo, S., & Jeong, T. (2024). Nerve signal transferring mechanism and mathematical modeling of artificial biological system design. Fractal and Fractional, 8(11), 648. 10.3390/fractalfract8110648

[44] Belykh, V. N., Osipov, G. V., Kuckländer, N., Blasius, B., & Kurths, J. (2004). Automatic control of phase synchronization in coupled complex oscillators. Physica D Nonlinear Phenomena, 200(1–2), 81–104. 10.1016/j.physd.2004.10.008

[45] Pikovsky, A., Rosenblum, M., & Kurths, J. (2000). PHASE SYNCHRONIZATION IN REGULAR AND CHAOTIC SYSTEMS. International Journal of Bifurcation and Chaos, 10(10), 2291–2305. 10.1142/s0218127400001481

[46] Kuznetsov, A., Seleznev, E., & Stankevich, N. (2012). Nonautonomous dynamics of coupled van der Pol oscillators in the regime of amplitude death. Communications in Nonlinear Science and Numerical Simulation, 17(9), 3740–3746. 10.1016/j.cnsns.2012.01.019

[47] Pikovsky, A., Rosenblum, M., Kurths, J., & Hilborn, R. C. (2002). Synchronization: a universal concept in nonlinear science. American Journal of Physics, 70(6), 655. 10.1119/1.1475332

[48] Elfouly, M. A., Fares, M. E., & Sohaly, M. A. (2025). Improved Tuberculosis Reinfection Dynamics Modeling with Delay Differential Equations. Research Square (Research Square). 10.21203/rs.3.rs-5656738/v1

[49] Elfouly, M. (2023). Hopf bifurcation and chaotic motion for Van der Pol model as two-delay differential equation in basal ganglia disorder. Brain Stimulation, 16(1), 305. 10.1016/j.brs.2023.01.553

[50] Al-Darabsah, I., Chen, L., Nicola, W., & Campbell, S. A. (2021). The impact of small time delays on the onset of oscillations and synchrony in brain networks. Frontiers in Systems Neuroscience, 15. 10.3389/fnsys.2021.688517

[51] Elfouly, M. A., Sohaly, M. A., & Fares, M. E. (2024). FitzHugh–Nagumo Model in Neutral Delay Differential Equation Representation. Research Square (Research Square). 10.21203/rs.3.rs-5048513/v1

[52] Elfouly, M. A., & Sohaly, M. A. (2021). Van der Pol Model in Two-Delay Differential Equation Representation. Research Square (Research Square). 10.21203/rs.3.rs-828074/v1

[53] Elfouly, M. A. (2024). Improved Mathematical Models of Parkinson’s Disease with Hopf Bifurcation and Huntington’s Disease with Chaos. Acta Biotheoretica, 72(3). 10.1007/s10441-024-09485-x

